# Investigation of cellular stress response related heat shock protein *hsp70*/Hsp70 and multixenobiotic transporter *abcb1* in Siberian freshwater amphipods upon cadmium exposure

**DOI:** 10.1101/626184

**Authors:** Marina V. Protopopova, Vasiliy V. Pavlichenko, Till Luckenbach

## Abstract

Induction of stress response genes *hsp70* and *abcb1* and Hsp70 protein by cadmium chloride (CdCl_2_) was explored in amphipod species with different stress adaptation strategies from the Lake Baikal area. Based on lethal concentrations (LC) the sensitivities to CdCl_2_ were ranked (24 hr LC50 – mg/L CdCl_2_): *Gammarus lacustris* (1.7) < *Eulimnogammarus cyaneus* (2.9) < *E. verrucosus* (8.3) < *E. vittatus* (18.2). Conjugated dienes indicating lipid peroxidation were significantly increased by 5 mg/L CdCl_2_ (24 hr exposure) only in *G. lacustris* and *E. cyaneus*. Upon treatment with 0.54 – 5.8 mg/L CdCl_2_ *hsp70* transcript levels were more increased in the toxicologically more sensitive species. Relating the exposure concentrations to LCx values revealed that across the species the increases of *hsp70* transcript levels were comparatively low (up to 2.6-fold) up to LC50; at higher LCx values *hsp70* induction was more pronounced (up to a 9.1-fold by 5 mg/L CdCl_2_ (≙LC70) in *E. cyaneus*). In contrast, *abcb1* inductions did not correspond with CdCl_2_ LCx values across species; *abcb1* induction was highest (4.7-fold) in *E. verrucosus* by 5.0 mg/L CdCl_2_ (≙LC45, 24 hr exposure). Induction of stress gene responses by lethal CdCl_2_ concentrations indicates that in the amphipods they are rather insensitive.

## Introduction

Cadmium is a non-essential heavy-metal that reaches the environment *via* various anthropogenic and natural sources and at low concentrations causes poisoning in humans and wildlife [1]. Toxic cadmium effects have often been attributed with increased levels of reactive oxygen species (ROS) and reactive nitrogen species (RNS) causing damages of biological macromolecules such as proteins [2]. Exposure to cadmium was found to lead to the induction of cellular stress response genes *hsp70* [3–12] and *abcb1* [13, 14] in a range of aquatic organisms. This induction of *hsp70* and *abcb1* by cadmium can be related to the increased abundance of damaged cellular macromolecules such as cellular membrane fragments or misfolded proteins [15, 16].

Lake Baikal in Eastern Siberia, the oldest, deepest and by volume largest lake in the world, is a biodiversity hotspot with high degrees of endemicity of species [17, 18]. Baikal’s water is generally highly pristine, however, the risk of water contamination by heavy metals is increasing especially through the Selenga river, which is the largest tributary of Lake Baikal and comprises almost half of the riverine inflow into the lake [19, 20]. Amphipods are a dominant taxon of the benthic communities of Lake Baikal and the more than 350 endemic species and subspecies represent 45.3% of all freshwater amphipod species of the world [21, 22]. The numerous phylogenetically closely related species featuring a range of adaptation strategies are interesting models for comparative studies [23]. In the here studied amphipod species constitutive expression levels of cellular stress response genes vary within an order of magnitude which could be related with differences across species regarding stress tolerance. Thus, constitutive *hsp70* levels relate to the species-specific differences in thermotolerance [24–26]. Furthermore, species-specific degrees of gene responses of *hsp70* and *abcb1* to organic compounds as humic substances [25] and phenanthrene [27] were found.

It is stated in many studies on the effects of cadmium on *hsp70*/Hsp70 levels that the test organisms were exposed to sublethal cadmium levels but the published molecular effect data are generally not related to the general level of stress caused by the cadmium concentration applied, such as lethal cadmium concentrations. It is thus unclear how sensitive the observed gene response to cadmium is.

We therefore here addressed the question at which stress levels cellular stress response genes *hsp70* and *abcb1* are induced upon exposure to CdCl_2_ in amphipod species with different cellular stress response capacities. The lethal concentrations (LCx 24h) of CdCl_2_ served as a measure for the cadmium-related stress levels. Transcript levels of *hsp70* and *abcb1* were quantified *via* quantitative polymerase chain reaction (qPCR) in RNA extracted from tissue of amphipods upon exposures of up to 24 hrs to different CdCl_2_ concentrations. The study was performed with four amphipod species that differ with regard to their ecological preferences and habitats and accordingly with their physiological and cellular adaptations to environmental conditions. Three of the species are littoral amphipods from Lake Baikal belonging to the *Eulinogammarus* genus; the Holarctic species *Gammarus lacustris* that occurs in waters connected to Lake Baikal but not in areas of the lake inhabited by endemic fauna was also examined.

## Materials and Methods

### Studied species and animal sampling

The experimental species were *Eulimnogammarus cyaneus* (Dyb., 1874), *Eulimnogammarus verrucosus* (Gerstf., 1858) and *Eulimnogammarus vittatus* (Dyb., 1874), endemic to Lake Baikal, and *Gammarus lacustris* (Sars, 1863) common in surface waters across the Holarctic. *Eulimnogammarus cyaneus* is most abundant along the shallow shoreline (mainly up to 1 m water depth), few individuals occur at water depths up to 20 m [28] at temperatures between 5-13°C [22]. Habitats of *E. vittatus* are at water depths of up to 30 m, the abundance peak is at 2–3 m depth [29]; the water temperatures in the habitat range between 9-13°C [22]. *Eulimnogammarus verrucosus* commonly occurs close to shore across Lake Baikal at water depths of less than 1 m to up to 10–15 m [29] at temperatures of 5-13°C [22]. *Gammarus lacustris* is common in shallow, eutrophic lakes with seasonal temperature fluctuations in the Lake Baikal region at water depths of 0-7m but the species is not found in the open Baikal [30]. Constitutive transcript/protein levels of heat shock protein 70 (*hsp70*/Hsp70) and transcript levels of ATP binding cassette (ABC) transporter 1 (*abcb1*) indicate the cellular stress response capacities of the species in the order: *Eulimnogammarus vittatus* ≈ *Eulimnogammarus verrucosus* < < *Eulimnogammarus cyaneus* < *Gammarus lacustris.* Indeed, the species-specific constitutive/induced *hsp70*/Hsp70 levels correlate with higher degrees of thermotolerance of *E. cyaneus* and *G. lacustris* compared to *E. verrucosus* [24–26, 31].

*Eulimnogammarus* specimens for the experiments were sampled at the Baikal shoreline close to Irkutsk State University’s biological station at the Bolshie Koty settlement (Southern Baikal) and *G. lacustris* was collected from a small shallow artificial lake close to Bolshie Koty (“Lake 14”, for specifications of the sampling points refer to [25]). Body lengths and weights of the animals used in the experiments were: *E. verrucosus* – 20-25 mm, 403 ± 95 mg; *E. vittatus* – 18-20 mm, 121 ± 16 mg; *E. cyaneus* – 9-14 mm, 17 ± 3 mg; *G. lacustris* – 14-18 mm, 80 ± 16 mg. Upon sampling the animals were brought to the lab in a cold box with water from the sampling sites. Animals were acclimated to lab conditions in aerated water in 2 L tanks at 6.5-7°C for 1-3 days prior to the experiments. The water used for maintaining the animals in the tanks and for the experiments was withdrawn from Lake Baikal with buckets from the pier next to the Biological Station in Bolshie Koty. Lake Baikal water instead of water from the pond where *G. lacustris* was collected was also used for *G. lacustris* exposures to avoid that differences in water contents (minerals, organic matter) affected the toxic effects of cadmium.

### 3.2. Acute toxicities of CdCl_2_

Acute toxicity of CdCl_2_ to amphipods was determined in 24 hr exposure experiments. Aqueous CdCl_2_ solutions for the experiments were set up in glass tanks with 2 L of well-aerated water along a control with clean water. The water temperature in the tanks was maintained at 6.5-7.0°C during the exposures. Twenty amphipods of each species were placed in separate tanks with eight different CdCl_2_ concentrations and a control. The CdCl_2_ concentration ranges were 0.5 – 8 mg/L for *G. lacustris* and *E. cyaneus* and 0.625-30 mg/L for *E. verrucosus* and *E. vittatus*. During the experiments, tanks were regularly checked for dead animals that were removed and recorded. After 24 hrs alive and dead animals in all tanks were counted. Exposure experiments were repeated two to three times with different CdCl_2_ concentrations that were chosen depending on the mortality rates determined for the already tested concentrations. Cadmium concentrations in water from a control and a CdCl_2_ exposure were determined with atomic absorption spectroscopy (AAS) at the Analytical Chemistry Department at the UFZ. The actual deviated from the nominal Cd concentration by 15 % (nominal: 10 CdCl_2_ mg/L; actual: 8.5 CdCl_2_ mg/L) and all nominal CdCl_2_ concentration values were adapted accordingly.

### 3.3. Measurements of conjugated diene (CD) levels

Levels of conjugated dienes (CD) were measured in amphipods upon exposure to 5 mg/L CdCl_2_ for 1, 6 and 24 hrs according to [32] with some modifications. Several entire animals were pooled to weights of 150 to 800 mg depending on the species and homogenized in a heptane/isopropyl alcohol mixture (1:1, v/v) using a Potter-Elvehjem tissue homogenizer, the extract was then filled in a glass tube and brought up to a volume of 4.5 ml with a heptane/isopropyl alcohol mixture. One ml of distilled water was added, the mix was vigorously shaken and incubated at 25°C for 30 min for phase separation. 0.5 ml of the heptane phase were mixed with ethanol in a 1:3 ratio (v/v) and the absorbance at 233 nm was measured on a SmartSpec Plus spectrophotometer (Bio-Rad) with extraction blanks used as references. An extinction coefficient of 2.52×10^4^ M^−1^cm^−1^ [33] was used to determine the amount of CD present. The data are reported as nmol g^−1^ wet weight.

### 3.4. CdCl_2_ exposures for investigating cellular stress responses

Two sets of exposure experiments were performed for examining cellular responses of the different amphipod species to CdCl_2_ exposure: 1) CdCl_2_ exposure concentrations were scaled to species-specific LC10 and LC50 values enabling to compare responses at equal stress levels. Respective CdCl_2_ exposure concentrations (LC10/LC50) were: *E. cyaneus* – 0.68/2.89 mg/L, *E. verrucosus* – 0.54/5.75 mg/L, *G. lacustris* – 0.59/1.7 mg/L; *E. vittatus* was only exposed to 5 mg/L corresponding to LC10. 2) Exposures were with all species at 1.7 (corresponding to the LC50 for *G. lacustris*) and 5.0 mg/L CdCl_2_ (corresponding to the LC90 for *G. lacustris*) enabling to determine species-specific responses to equal concentrations. CdCl_2_ exposures were performed in aerated 2 L tanks (n=5/concentration) with Baikal water at 7°C. Specimens were sampled after 1, 6, and 24 hr exposures, frozen and stored in liquid nitrogen for extraction of total protein and diene conjugates or placed in QIAzol Lysis Reagent (Qiagen, Hilden, Germany) and frozen in liquid nitrogen for RNA isolation. Control animals were kept in CdCl_2_ uncontaminated water but otherwise at equal temperature and aeration conditions as CdCl_2_ exposed animals. One whole specimen of *E. verrucosus* or a pool of several whole animals of the other species equalling 200 to 500 mg of tissue were used for the further analyses. All exposure experiments were repeated at least five times and all samples were assessed in triplicate.

### 3.5. Isolation of total RNA and cDNA synthesis

Isolation of total RNA and cDNA synthesis were performed as described in [27]. Tissue preserved in QIAzol Lysis Reagent was homogenized with a MM400 homogenizer (Retsch, Haan, Germany) followed by a phenol-guanidine-chloroform based extraction of total RNA. To improve the separation of the organic and aqueous phases MaXtract gel (Qiagen) was used according to the manufacturer. Total RNA was purified from the obtained aqueous phase with the miRNeasy kit with DNase treatment using a QIAcube instrument (Qiagen). One μg of total RNA was reverse-transcribed to single-stranded cDNA using Oligo(dT)18 primer (Fermentas, USA, #SO131) and H Minus Reverse Transcriptase system (Fermentas, USA, #EP0452) following the manufacturer’s instructions.

### 3.6. Quantitative real-time PCR analysis

Gene expression levels were analysed with quantitative polymerase chain reaction (qPCR) using a StepOnePlus Real-Time PCR System (Applied Biosystems) as described previously [25]. The amplification was performed in a final volume of 12.5 μl using the SensiMix SYBR Low-ROX Kit (Bioline) with each primer at 200 nM. Primer sequences for qPCR of *hsp70* and *abcb1* and housekeeping genes *β-actin, gapdh* and *ef1-a* were from [25]; qPCR conditions were according to [25, 27]: 94 °C for 4 min; 35 (for *hsp70* and references genes) and 45 cycles (for *abcb1*) with 95 °C for 15 sec, 60-62 °C for 15 sec and 72 °C for 15 sec followed to fluorescent measurement step with 78-79°C (close to the melting temperature of the PCR products) for 10 sec. The extra step for fluorescence measurement was added to avoid detection of unspecific amplification products and primer dimers. Relative expression levels of *hsp70* and *abcb1* genes were calculated with the comparative ΔΔCt method [34] using efficiency corrected calculation models and Best Keeper levels based on the geometric mean of the Ct values of the housekeeping genes [35–37].

### 3.7. Western-blotting

Isolation of total protein from amphipod tissue and Western-blotting were performed as described in [25]. Tissues were homogenized in 0.1 M Tris buffer (pH=7.6) containing 1.5 mg phenylmethylsulfonyl fluoride (PMSF) and 1% protease inhibitor cocktail (Amresco, USA); proteins were then subjected to denaturation in Tris-HCl buffer (0.25 M Tris-HCl (pH = 6.8), 5% sodium dodecyl sulphate (SDS), 0.5 mM ethylenediamine tetraacetic acid (EDTA), 10% glycerol, 5% ß-mercaptoethanol and 0.03% Bromophenol Blue) at 95 °C for 5 min. Total protein concentrations were determined following [38]. Equal amounts of total protein per sample were separated by SDS gel electrophoresis on a 12.5% polyacrylamide gel [39] using a Mini-PROTEAN II Electrophoretic Cell (Bio-Rad, USA). Wet-transfer of the proteins from the gel to a polyvinylidene difluoride transfer membrane (GE Healthcare, UK) was according to [40] with minor modifications. Hsp70 protein was labelled with anti-HSP70 (Sigma-Aldrich, # H9776) at 1:3000, which detects both the inducible and constitutive forms of HSP70, and with secondary antibodies conjugated with alkaline phosphatase (Anti-Mouse IgG:AP Conj., Stressgen # SAB-101) at 1:1000. Anti-actin antibody (Sigma-Aldrich, # A2668) dissolved 1:400 and secondary antibodies (Anti-Rabbit IgG, Sigma, # A9919) dissolved 1:1000 were used for actin labelling. Sixty ng of bovine Hsp70 (Sigma-Aldrich, # H9776) and 60 ng of bovine actin (Sigma-Aldrich, A3653) were used as positive controls. Bands were visualized with 0.4 mM BCIP (5-bromo-4-chloro-3-indolyl phosphate) and 0.4 mM nBT (nitroblue tetrazolium). The Gel Explorer software package (DNAtechnology, Russia) was used for semi-quantitative measurements of Hsp70 and actin levels on the membranes and Hsp70 levels were normalized relative to actin for each sample.

### 3.8. Data analysis and statistics

Regressions of concentration-mortality relationships were calculated by fitting mortalities at the respective concentrations to the non-linear HILL model:

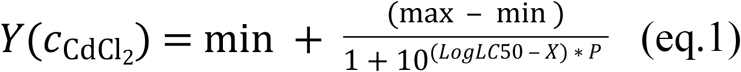

where

Y: is the mortality at the CdCl_2_ concentration *c*
X: is the logarithm of the CdCl_2_ concentration
min: is the minimum percentage mortality (control, constrained to 0)
max: is the maximal percentage mortality (constrained to 100)
P: is the shape parameter
LC50: is the constant describing the CdCl_2_ concentration causing 50% mortality

Regressions of *hsp70* transcript/Hsp70 protein levels at respective CdCl_2_ concentrations expressed as LCx values were calculated by fitting the observations at the respective concentrations to the Gaussian distribution model:

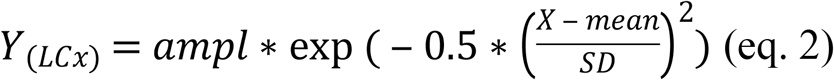

where

Y: is the gene expression level at LCx (lethal concentration for x% of individuals)
X: is LCx
ampl: (amplitude) is the height of the center of the distribution in Y units
SD: is a measure of the width of the distribution (in LCx)

Data for qPCR, Western-blot and lipid peroxidation analyses were found to satisfy the assumptions of equal variance and normality and parametric statistics were used. For pairwise comparisons of *abcb1* and *hsp70*/Hsp70 levels in treatments and respective controls at each time point the t-test was applied. Multiple comparisons of several treatments with one respective control were done with two-way ANOVA and Dunnett’s test. To analyse the significance of observed changes in each time-point, pair-wise comparisons with the respective control were done. The results of multiple pair-wise comparisons were done using Steel-Dwass method for all pairs’ comparison. Differences were regarded as significant if *P* < 0.05. Regressions were calculated with Graphpad Prism version 7. Statistical analyses were performed with JMP version 10.0 (SAS Institute, Cary, NC).

## Results

### 4.1. Lethal CdCl_2_ concentrations

Acutely toxic CdCl_2_ concentrations were across the different amphipod species from < 1 to < 100 mg/L (Fig. 1, Table 1). The range of LCx values calculated from CdCl_2_ concentration – lethality relationships were over an order of magnitude across the species. LCx values were highest for *E. vittatus* indicating the lowest sensitivity of this species to CdCl_2_. The order of the species with regard to their sensitivities to CdCl_2_ was: *G. lacustris < E. cyaneus < E. verrucosus < E. vittatus* (Fig. 1, Table 1).

**Table 1.** Lethal concentration values (LCx; x: % lethality) for CdCl_2_ (mg/L) and curve parameters for the different amphipod species determined with the HILL model (eq. 1, see Fig. 1). CI: confidence interval, R^2^: curve fit

|  |  | *E. vittatus* | *E. verrucosus* | *E. cyaneus* | *G. lacustris* |
| --- | --- | --- | --- | --- | --- |
| LC<sub>x</sub> | 10 | 4.47 | 0.54 | 0.67 | 0.59 |
|  | 25 | 8.98 | 1.76 | 1.38 | 1.01 |
|  | 30 | 10.63 | 2.31 | 1.64 | 1.14 |
|  | 45 | 16.07 | 4.65 | 2.52 | 1.55 |
|  | 50 | 18.05 | 5.72 | 2.89 | 1.71 |
|  | 95% CI | 12.4-26.6 | 3.8-8.7 | 2.1-4.0 | 1.3-2.4 |
|  | 70 | 31.05 | 14.31 | 5.05 | 2.59 |
|  | 75 | 36.74 | 18.76 | 5.96 | 2.91 |
|  | 90 | 73.81 | 60.81 | 12.22 | 4.98 |
|  | HillSlope | 0.34 | 0.20 | 0.34 | 0.53 |
|  | 95% CI | 0.7 - 2.4 | 0.5 - 1.4 | 0.7 - 2.3 | 0.8 - 3.3 |
|  | R<sup>2</sup> | 0.88 | 0.83 | 0.87 | 0.88 |

**Figure 1.**
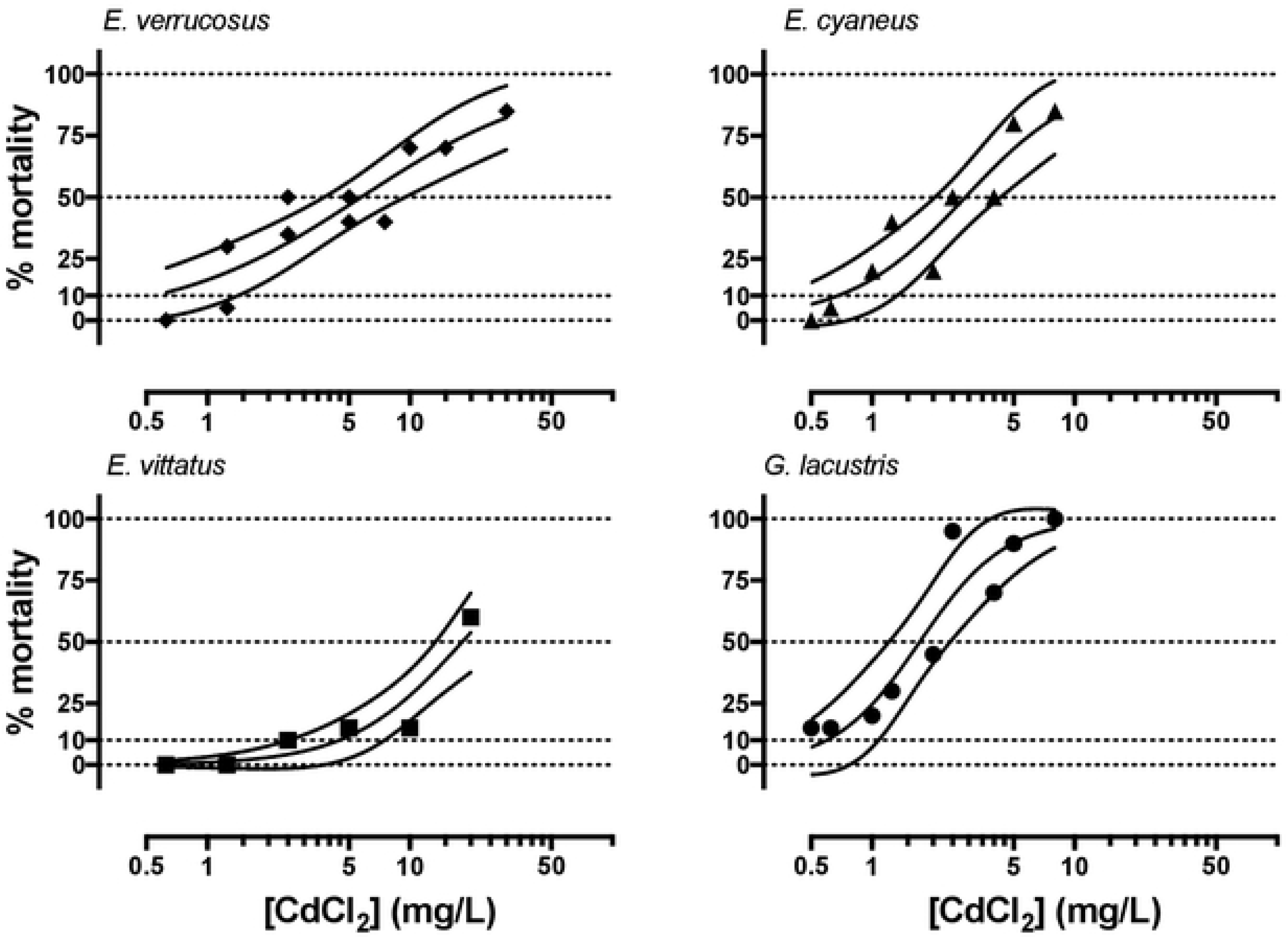
Concentration-mortality relationships for CdCl_2_ in the different amphipod species. Symbols represent the observed % mortality at a certain CdCl_2_ concentration in a single treatment. Solid lines represent the regression curves fitted with the HILL model (eq. 1) and dashed lines the 95% confidence intervals.

### 4.2. Conjugated diene (CD) levels

After 1 and 6 hr exposures to 5 mg/L CdCl_2_ no significant changes in CD levels were seen in any of the examined species. Significant increases of CD levels by about 58% and 47% occurred only in *E. cyaneus* and *G. lacustris*, respectively, after 24 hr exposures (Fig. 2). The CdCl_2_ concentration of 5 mg/L corresponds to LC70 and LC90 for *E. cyaneus* and *G. lacustris*, respectively. For *E. vittatus* and *E. verrucosus* this CdCl_2_ concentration is equivalent to LC10 and LC45, respectively (Fig. 1, Table 1).

**Figure 2.**
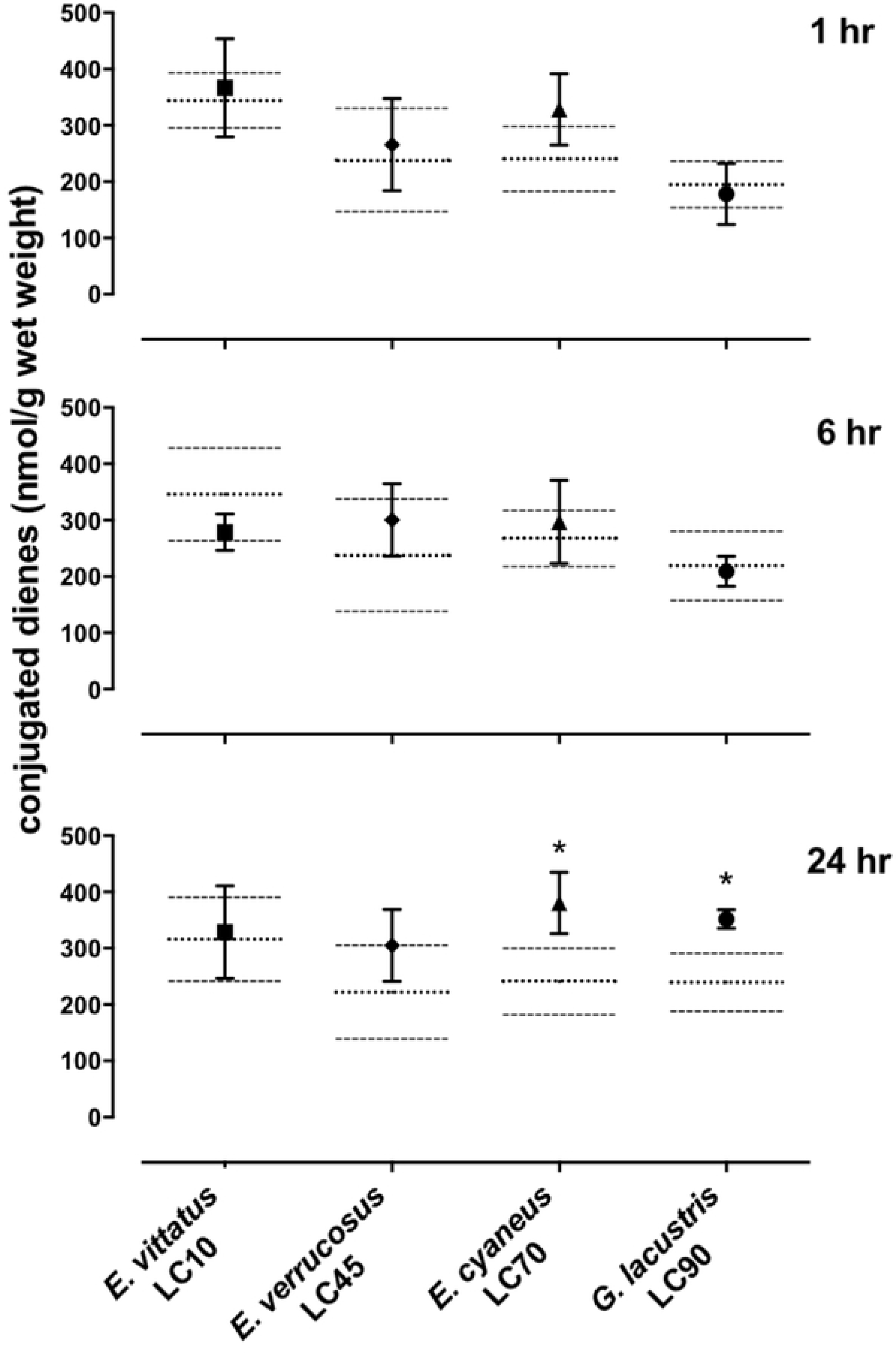
Levels of conjugated dienes (CD; nmol g^−1^ wet weight) in the different amphipod species upon exposure to 5 mg/L CdCl_2_ that correspond to the species-specific LCx values given under each species name. Depicted are means (symbols) and standard deviations (bars) in the CdCl_2_ exposed animals. The control levels are shown as dotted (mean) and dashed (standard deviations) lines. Significant differences between treatments and respective controls are indicated by * (p< 0.05).

### 4.3. Heat shock protein 70 transcript/protein levels

*Hsp70* transcript levels across the amphipod species were upon exposures to CdCl_2_ significantly (p < 0.05) increased by between 1.4 - (*E. cyaneus* at LC30 for 24 hrs) to up to 9.1-fold (*E. cyaneus* at LC70 for 24 hrs) (Fig. 3a, b). With longer exposure times to CdCl_2_ changes in *hsp70* transcript levels were in all amphipod species generally more pronounced: After 1 hr exposures *hsp70* levels were significantly (p < 0.05) increased in two cases (*E. cyaneus* at LC30, *G. lacustris* at LC90), after 6 hr exposures in four cases (*E. cyaneus* at LC30, *E. verrucosus* and *G. lacustris* at LC50, *G. lacustris* at LC90) and after 24 hr exposures in nine cases (*E. vittatus* at LC10, *G. lacustris* at LC10, LC50 and LC90, *E. verrucosus* at LC25, LC45 and LC50, *E. cyaneus* at LC30 and LC70; Fig. 3a). Although significant changes in *hsp70* transcript levels were also seen with CdCl_2_ at lower LCx, changes were more pronounced at higher LCx levels and could be described with a regression curve over all data following a Gaussian shape with an overall maximum *hsp70* transcript level at LC70 followed by a decrease (Fig. 3b).

**Figure 3.**
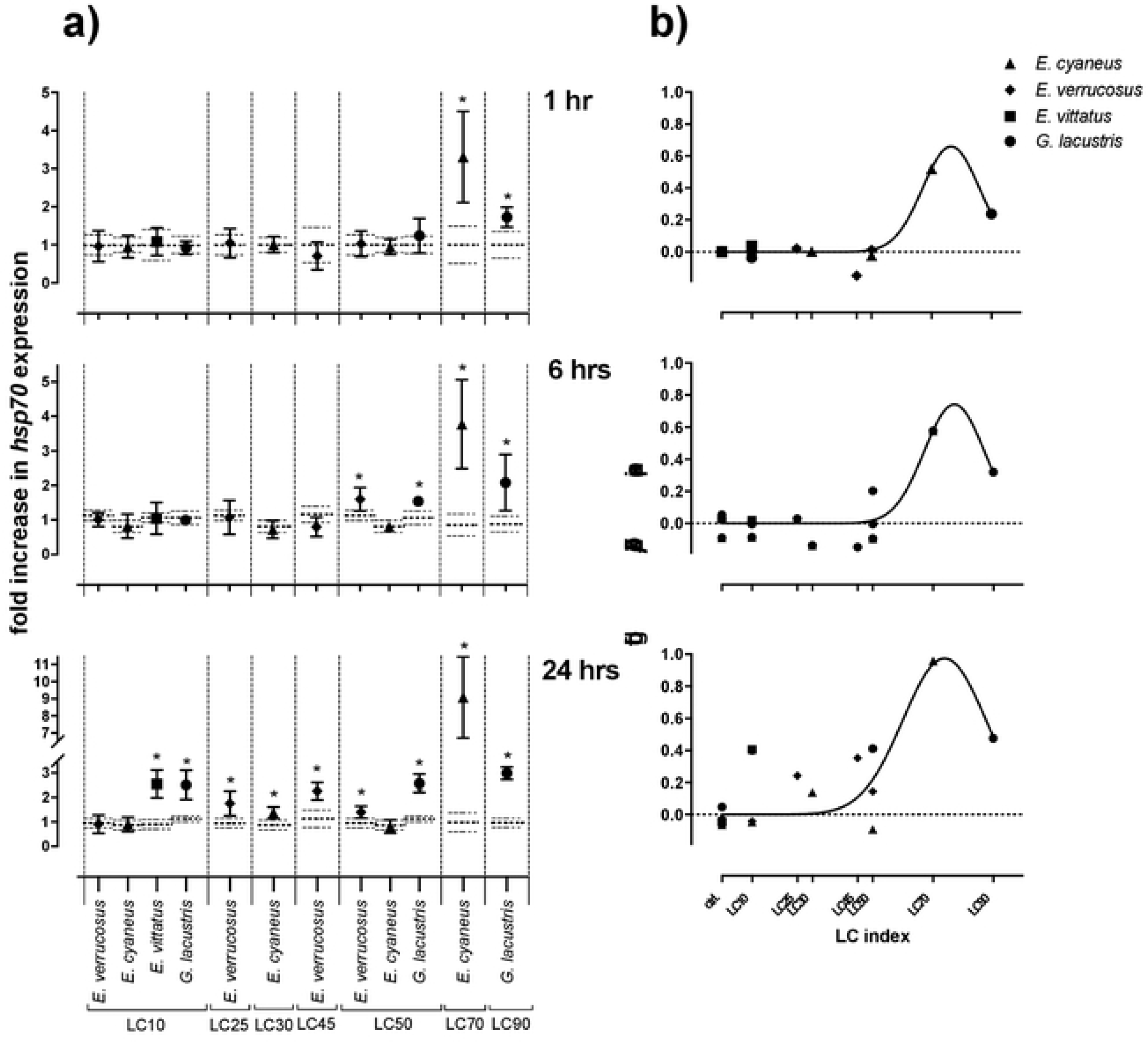
*Hsp70* transcript levels in tissue of the different amphipod species upon exposure to CdCl_2_ for 1, 6 and 24 hrs. CdCl_2_ concentrations were scaled to the species-specific LCx values (refer to Table 1 for the LCx equivalences in mg/L). a) Depicted are mean *hsp 70* expression levels (symbols) and standard deviations (bars) in CdCl_2_ exposed animals. Expression levels in controls are shown as dotted (mean) and dashed (standard deviations) lines. Significant differences between treatments and respective controls are indicated by * (p< 0.05). b) The *hsp 70* expression level data of all species were combined and are depicted in relation to the correspondent species-specific LCx value. The regression line was fitted over all data with the Gaussian distribution model (eq. 2).

Hsp70 protein levels across amphipod species from CdCl_2_ treatments also followed a Gaussian function when expression data were related to the respective LCx values (Fig. 4). Significantly (p < 0.05) increased Hsp70 protein levels were seen in *E. cyaneus* at 1.8- and 1.7-fold, respectively, after 1 and 6 hr exposures to 5 mg/l CdCl_2_ corresponding to the species-specific LC70; and in *E. verrucosus* after 24 hrs exposure with a 2.6-fold increased protein level above the control. For this species 5 mg/l CdCl_2_ corresponded to the LC45. In *E. vittatus* and *G. lacustris* exposed to the species-specific LC10 and LC90, respectively, Hsp70 protein levels were not significantly different from the respective controls (Fig. 4).

**Figure 4.**
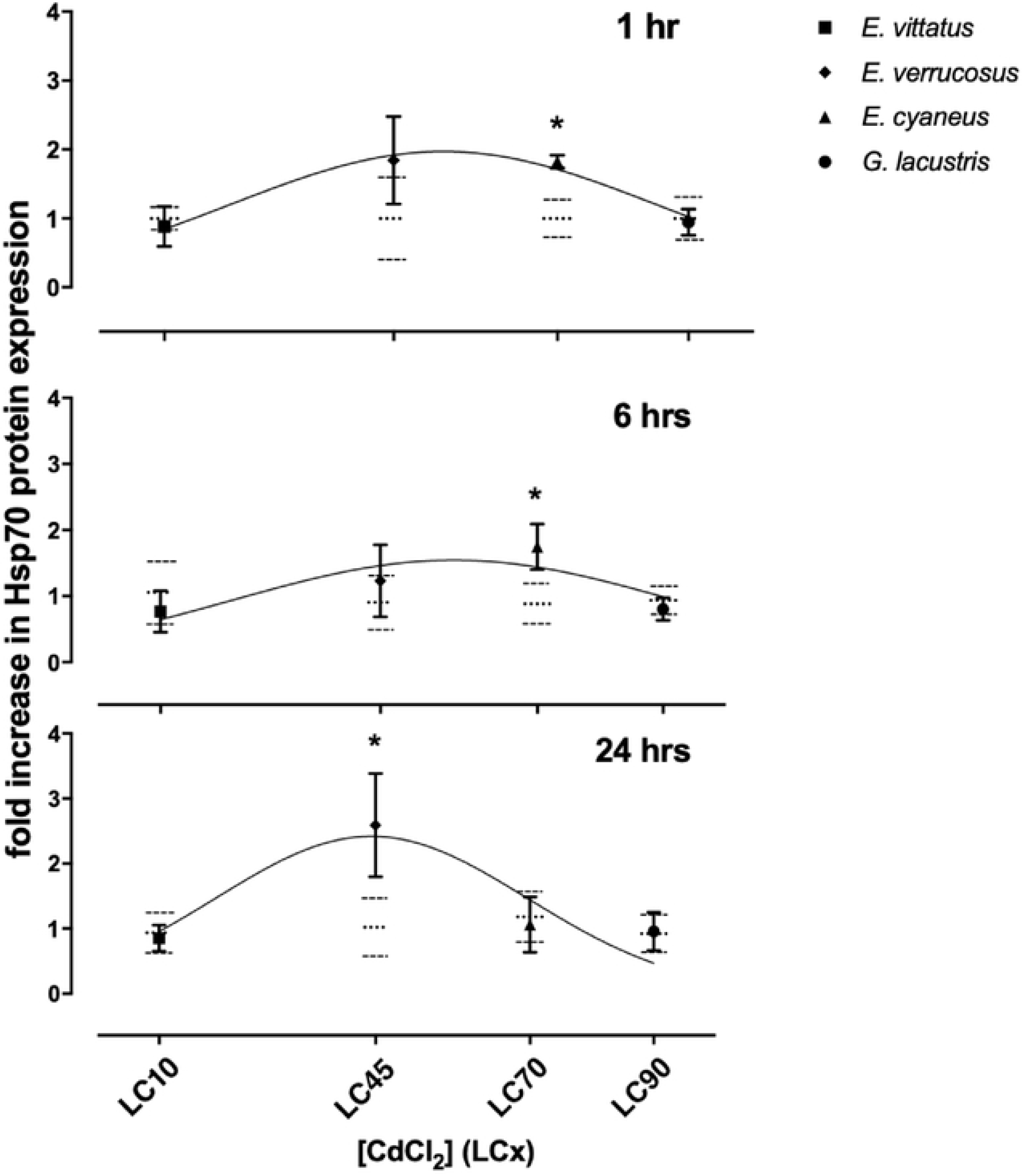
Hsp70 protein levels in tissue of the different amphipod species upon exposure to 5 mg/L CdCl_2_ for 1, 6 and 24 hrs. CdCl_2_ concentrations were scaled to the species-specific LCx values. Depicted are mean Hsp70 expression levels (symbols) and standard deviations (bars) in CdCl_2_ exposed animals. Expression levels in controls are shown as dotted (mean) and dashed (standard deviations) lines. Significant differences between treatments and respective controls are indicated by * (p< 0.05). The regression line was fitted with the Gaussian distribution model (eq. 2).

### 4.4. Abcb1 transcript levels

Upon exposures of amphipods to CdCl_2_ *abcb1* transcript levels were significantly (p < 0.05) increased by between 2.2- (*E. cyaneus* at LC50 for 24 hrs) to up to 4.7-fold (*E. verrucosus* at LC45 for 24 hrs). Over the different exposure times the number of cases of *abcb1* increases only slightly changed. Thus, *abcb1* transcript levels were increased in three cases at 1 hr exposures (*E. verrucosus* at LC25, *E. cyaneus* at LC70, *G. lacustris* at LC90), in two cases at 6 hr exposures (*E. cyaneus* at LC50, *G. lacustris* at LC90) and in four cases at 24 hr exposures (*E. verrucosus* at LC45, *E. cyaneus* at LC50, *G. lacustris* at LC90, *E. vittatus* at LC10). Significant *abcb1* transcript increases at all three time points occurred only in *G. lacustris* at LC90; in *E. cyaneus* at LC50 *abcb1* transcripts were increased at two time points. In the other cases, significant *abcb1* transcript increases were seen only at one time point. Changes in *abcb1* transcript levels did not seem to depend on CdCl_2_ concentration / LCx value; the significant changes were seen across different LCx values and were not generally induced at higher or lower LCx values (Fig. 5).

**Figure 5.**
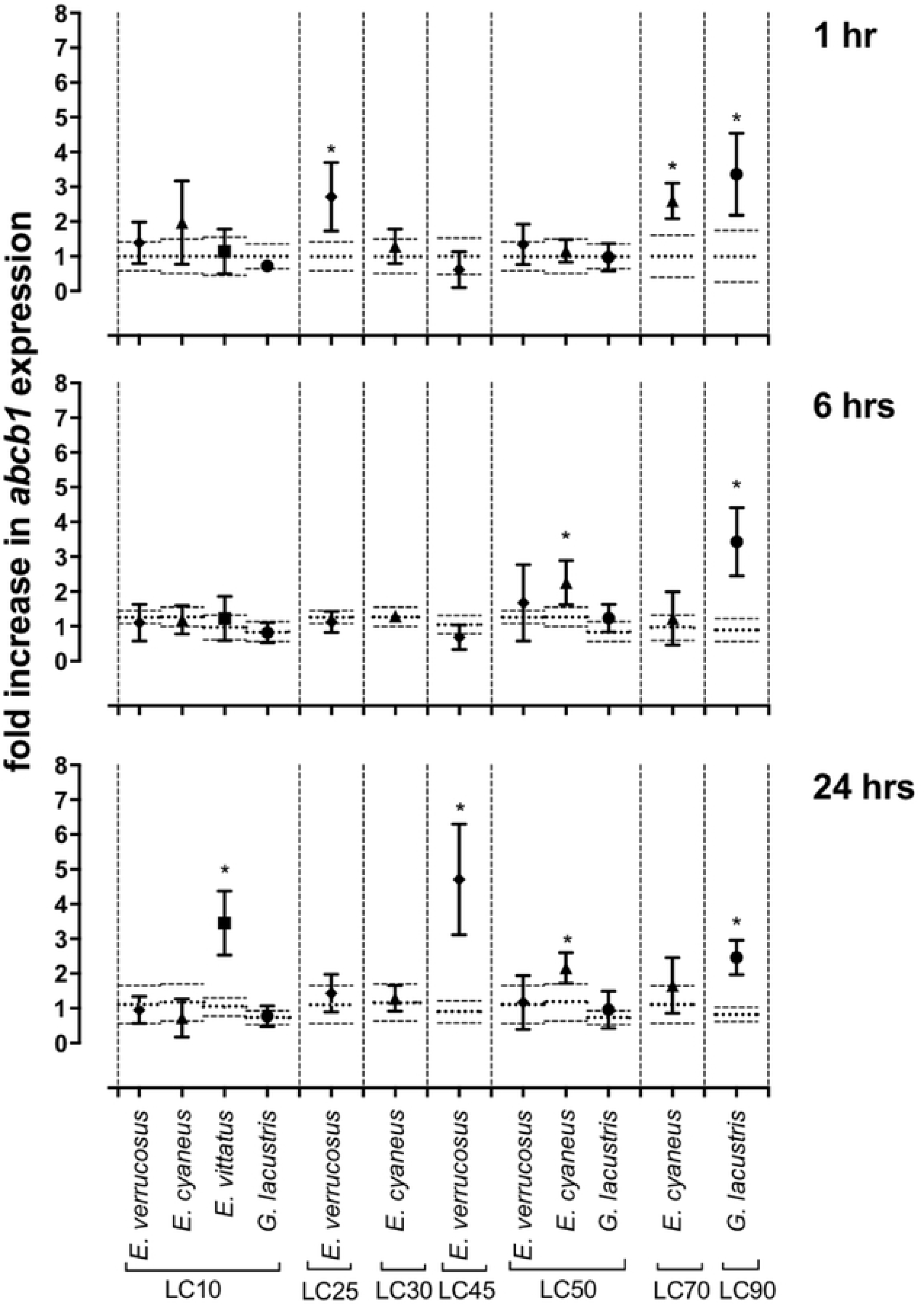
Relative *abcb1* levels in amphipods upon exposure to CdCl_2_ for 1, 6 and 24 hrs. CdCl_2_ concentrations were scaled to the species-specific LCx values. Depicted are mean *abcb1* levels (symbols) and standard deviations (bars) in CdCl_2_ exposed animals. Expression levels in controls are shown as dotted (mean) and dashed (standard deviations) lines. Significant differences between treatments and respective controls are indicated by * (p< 0.05).

## Discussion

The four examined amphipod species showed clear differences regarding toxic sensitivities to CdCl_2_ that became evident by LC50 values ranging over one order of magnitude (Table 1). Along those lines the levels of conjugated dienes, indicating cadmium effects on the molecular level, were at equal CdCl_2_ exposure concentrations only increased in the toxicologically more sensitive species *E. cyaneus* and *G. lacustris* (Fig. 2). Conjugated dienes are primary products of lipid peroxidation resulting from oxidative stress [41, 42], a main cause of the toxic action of cadmium [43, 44]. This effect of CdCl_2_ on gammarids was indicated in a previous study where it was found that CdCl_2_ exposure leads to the inhibition of common antioxidant enzymes in gammarids [45]. Differences in internal cadmium levels as a consequence of different cadmium uptake rates can, to a certain extent, be seen as a reason for the differences in toxicological sensitivities of the different species. Body size has previously been shown to be an important determinant of cadmium uptake by aquatic insects [46]. Along those lines, cadmium uptake rates during CdCl_2_ exposures were higher in the concurrently toxicologically more sensitive *E. cyaneus* than in *E. verrucosus* that differed by 10- to 50-fold in freshweight [47]. In the present study, *G. lacustris* individuals that were similar in size to *E. cyaneus* individuals showed toxic sensitivities that were accordingly in the same range as for *E. cyaneus* (Table 1). *Eulimnogammarus vittatus*, however, was from the examined species least sensitive to CdCl_2_ although the body sizes of the individuals were clearly below those of *E. verrucosus* and it can be assumed that the cadmium uptake rate was accordingly higher. It was previously found that the occurrence of toxic effects by CdCl_2_ in *E. verrucosus* was in addition to a in comparison to *E. cyaneus* lower cadmium uptake rate further depressed by a reaction to CdCl_2_ exposure with metabolic depression [47]. This was not examined here but it seems conceivable that the in comparison low toxic sensitivity of *E. vittatus* to CdCl_2_ may be due to particularly pronounced metabolic depression as a reaction to CdCl_2_ exposure.

Although metabolic depression by cadmium may have decreased activity on a physiological level it appeared to have no effect on the transcription activity. Thus, *hsp70* was significantly induced in *E. verrucosus* that previously showed a metabolic depression response to CdCl_2_ exposure but not in *E. cyaneus* that does not respond to CdCl_2_ with metabolic depression [47] in the 6 and 24 hr CdCl_2_ treatments at LC50 (Fig. 3). Furthermore, a significant *abcb1* increase was seen in *E. verrucosus* at LC25 but not in *E. cyaneus* at LC30 after 1 hr exposure to CdCl_2_ and in *E. vittatus* but no other species at LC10 after 24 hrs exposure to CdCl_2_ (Fig 5).

The CdCl_2_-dependent *hsp70* transcript and Hsp70 protein response levels could across the species be related to the respective CdCl_2_–related stress levels. When plotting all *hsp70* transcript or Hsp70 protein levels across species against the respective species-specific LCx values the values showed an increase to a maximum followed by a decrease at higher LCx values (Figs. 3, 4). When assuming that the increasing LCx values represent increasing degrees of stress this course represents stress-level dependent expression degrees that for Hsp70 has been described before: The Hsp70 titre increase in the “compensation phase” is followed by a Hsp70 titre decrease in the “non-compensation phase” at higher toxicity levels when the ability of the cells to react to the toxicant is increasingly degraded [48]. The applied CdCl_2_ concentration represented different degrees of stress for the different amphipod species; at this stage it is assumed but it cannot be definitely asserted that the *hsp70* transcript/Hsp70 protein responses are equal in the different amphipod species at corresponding stress levels.

Provided that across the gammarid species equal internal, toxicologically relevant cadmium concentrations cause equal levels of toxic effects, as was indicated earlier [47], the LCx values can be regarded as approximate measures of the respective internal cadmium concentrations. Hence, across the species the degrees of *hsp70* transcript and Hsp70 protein induction appear to be approximately equal at corresponding internal cadmium levels.

Induction of *abcb1* transcripts tends to depend on LCx levels and time of exposure, i.e., internal cadmium levels. Increased *abcb1* levels were more seen at higher CdCl_2_ LCx values and occurred at lower LCx levels after longer times of exposure (Fig. 5). However, across species *abcb1* induction levels are not uniform at equal LCx levels. Thus, after 24 hrs exposure to CdCl_2_ at LC10 *abcb1* levels were only significantly increased in *E. vittatus* but not in *E. verrucosus*, *E. cyaneus* and *G. lacustris*; after 6 and 24 hrs exposure to CdCl_2_ at LC50 significant *abcb1* increases were only seen in *E. cyaneus* but not in *E. verrucosus* and *G. lacustris* (Fig. 5).

The differences in sensitivity of the *hsp70* and *abcb1* transcript responses among the gammarid species may confirm that induction of the two genes is *via* different pathways. Misfolded proteins that are damaged e.g. by the impact of cadmium-induced ROS [15, 16] trigger the induction of the *hsp70* gene as, according to the chaperone titration model [49], heat shock factor (Hst1) is released when Hsps bind to the misfolded proteins thus initiating *hsp* transcription. Induction of *abcb1* appears to be triggered *via* another pathway involving the transcription factor NF-κB [15, 16].

Both Hsp70 and Abcb1 proteins act as protective proteins against cadmium toxicity: Hsp70 as chaperone preserves the structure of intact proteins and re-folds damaged proteins; Abcb1, a cellular efflux transporter, appears to efflux cell membrane components, such as ceramides, that result from membrane damage by ROS and can induce apoptosis [15, 16]. It was found earlier that *E. cyaneus* that shows higher constitutive and induced levels of Hsp70 than *E. verrucosus* [24, 25] detoxifies a higher proportion of cadmium by binding it to proteins of the biologically detoxified heat stable protein fraction (BDF) [47]. The in comparison high constitutive Hsp70 titres in *Gammarus lacustris* [25] may indicate that also in this species a comparatively high level of the internal cadmium is bound to the BDF. However, when considering that the toxicologically more sensitive gammarid species have higher titers of cadmium detoxifying proteins and that the transcripts of both here examined proteins are induced by cadmium only at comparatively high stress levels, it seems obvious that overall the detoxifying effect by those cellular stress response proteins on cadmium is only minor. Major determinants of the cadmium sensitivity of the species appear to be body size and in relation to that the cadmium uptake rate and one additional parameter that could be the degree of metabolic depression caused by cadmium.

## Summary and Conclusions

In this study cadmium induced *hsp70*/Hsp70 and *abcb1* levels in gammarids exposed to CdCl_2_ were for the first time phenotypically anchored, i.e., exposure concentrations of treatments for transcript measurements were equated with the respective lethal concentrations (LCx). When gene/protein responses were related to LCx values the four gammarid species examined showed a relatively uniform *hsp70*/Hsp70 response with pronounced responses in CdCl_2_ treatments at higher LCx and in lower LCx treatments after longer times of exposure to CdCl_2_ (Figs. 3, 4). For *abcb1*, responses to CdCl_2_ exposure when related to LCx levels were not uniform across the species; however, *abcb1* was overall also induced by CdCl_2_ in the lethal concentration range (Fig. 5). Induction of *hsp70*/Hsp70 and *abcb1* responses by CdCl_2_ in the lethal concentration range in the gammarids indicates that changes in transcript levels of those genes are rather insensitive markers for cadmium stress.

## Acknowledgements

We would like to thank Rolf Altenburger, Andreas Schüttler, Daria Bedulina and Maxim Timofeyev for their input and discussions related to the study; Sibylle Mothes for Cd analyses of water samples; and Stephan Schreiber for his help with the QIAcube at the Fraunhofer Institute for Cell Therapy and Immunology IZI, Leipzig. This study was financially supported by scholarships from the Deutscher Akademischer Austauschdienst (DAAD) and the Russian Ministry of Education and Science (“Mikhail Lomonosov” Programme) (M.P., V.P.), by an international DAAD scholarship (M.P., V.P.), by an Erasmus Mundus (MULTIC II) scholarship (V.P.), the project of Goszadaniye of Siberian Institute of Plant Physiology and Biochemistry (No 0343-2017-0006; M.P., V.P.) and by the bilateral funding programs “Helmholtz-Russia Joint Research Groups” (HRJRG) from the Helmholtz Association and the Russian Foundation for Basic Research (RFBR) (LaBeglo project HRJRG-221; M.P., V.P., TL) and „Helmholtz-RSF Joint Research Groups” from the Helmholtz Association and the Russian Science Foundation (RSF) (LaBeglo2/RSF 18-44-06201; T.L.). The funders had no role in study design, data collection and analysis, decision to publish, or preparation of the manuscript.

## Data accessibility statement

All raw data can be obtained from the authors upon request.

